# GS-Preprocess: Containerized GUIDE-seq Data Analysis Tools with Diverse Sequencer Compatibility

**DOI:** 10.1101/2020.01.26.914861

**Authors:** Tomás C. Rodríguez, Henry E. Pratt, PengPeng Liu, Nadia Amrani, Lihua Julie Zhu

## Abstract

RNA-guided nucleases (e.g. CRISPR-Cas) are used in a breadth of clinical and basic scientific subfields for the investigation or modification of biological processes. While these modern platforms for site-specific DNA cleavage are highly accurate, some applications (e.g. gene editing therapeutics) cannot tolerate DNA breaks at off-target sites, even at low levels. Thus, it is critically important to determine the genome-wide targeting profile of candidate RNA-guided nucleases prior to use. GUIDE-seq is a high-quality, easy-to-execute molecular method that detects and quantifies off-target cleavage. However, this method may remain costly or inaccessible to some researchers due to its library sequencing and analysis protocols, which require a MiSeq platform that must be preprogramed for non-standard output. Here, we present GS-Preprocess, an open-source containerized software that can use standard raw data output (BCL file format) from any Illumina sequencer to create input for the Bioconductor GUIDEseq off-target profiling package. Single-command GS-Preprocess performs FASTQ demultiplexing, adapter trimming, alignment, and UMI reference construction, improving the ease and accessibility of the GUIDE-seq method for a wide range of researchers.

## Introduction

CRISPR-Cas nucleases are an increasingly practical tool for the interrogation of nuclear events, largely due to their programmability and the wide variety of low-cost constructs available for clinical and laboratory applications. However, target genomic sites are still limited to those that have a corresponding guide RNA (gRNA) verified to cause little or no off-target cleavage by Cas nucleases (Fu et al. 2013; Doench et al. 2016; Anderson et al. 2018). While computational techniques provide a free and expedient means of predicting promiscuous gRNA activity (Bae, Park, and Kim 2014; Listgarten et al. 2018), molecular validation is indispensable for the detection of off-targeting in a cellular context—especially those containing high spacer mismatches ignored by computational models (Tsai et al. 2015).

GUIDE-seq (<u>g</u>enome-wide, <u>u</u>nbiased <u>i</u>dentification of <u>d</u>ouble-stranded breaks enabled by <u>seq</u>uencing) is a high-fidelity method for establishing the targeting profile of novel gRNA designs with high sensitivity and specificity (Tsai et al. 2015). GUIDE-Seq relies on the incorporation of a small double-stranded oligodeoxynucleotide (dsODN) wherever there is an RNA-guided Cas-induced break in the genome. dsODN integration sites are then mapped in the genome using unbiased amplification and next generation sequencing (NGS). Relatively low read-depth requirements combined with growing ease of performing highly-multiplexed NGS make GUIDE-seq a sensible addition to CRISPR experiments. Yet, despite being relatively straightforward to carry out, GUIDE-seq could be more widely used with higher sample plexity and expanded compatibility with Illumina sequencers. Current protocols also require users to preinstall third-party software on a MiSeq to generate output data in a non-standard format, and to use a YAML manifest that is not immediately modifiable by commonplace, user-friendly programs like Microsoft Excel or Apple Numbers (https://github.com/aryeelab/guideseq#miseq).

Here, we present GS-Preprocess, an open-source package that partially subverts existing equipment and computation skill requirements to sequence and analyze GUIDE-seq libraries. GS-Preprocess uses standard raw output from *any* Illumina sequencer to generate input data compatible with the GUIDEseq Bioconductor (BC) package in R (Zhu et al. 2017). We chose to work with GUIDEseq BC because it can be easily paired and containerized with GS-Preprocess due to its BC-bundled dependencies and single-line execution.

## Results

### Implementation and Functionality

In a one-line, six-argument command, GS-Preprocess creates de-multiplexed aligned sequences in BAM file format, a tab-separated Unique Molecular Index (UMI) reference, and gRNA sequences in a fasta formatted file (Fig. 1a), all of which can be fed directly into the GUIDEseq BC package. Using the well-documented bcl2fastq software (Illumina), sequencer raw data output in standard BCL format is converted to demultiplexed FASTQ files and UMI-containing Index reads (Fig. 1b, adapted from Tsai et al. 2015). Fig. 1a outlines the location of each library adapter component with respect to Read/Index. A custom python script mines Index reads to build a UMI reference. Adapters are removed from demultiplexed FASTQs using the command-line tool cutadapt (Martin 2011). Trimmed FASTQs are then aligned to a user-provided reference genome with the BWA-MEM algorithm (Li 2013) and compressed into BAM format with samtools. (Li et al. 2009) To facilitate usability and reproducibility, we containerized GS-Preprocess, GUIDEseq BC package and all dependencies on the docker platform (umasstr/gsp). Detailed instructions can be found at github.com/umasstr/GS-Preprocess in the README.md file. Instructions for optional merging of lane-separated data or technical replicates are also provided.

**Figure 1:**
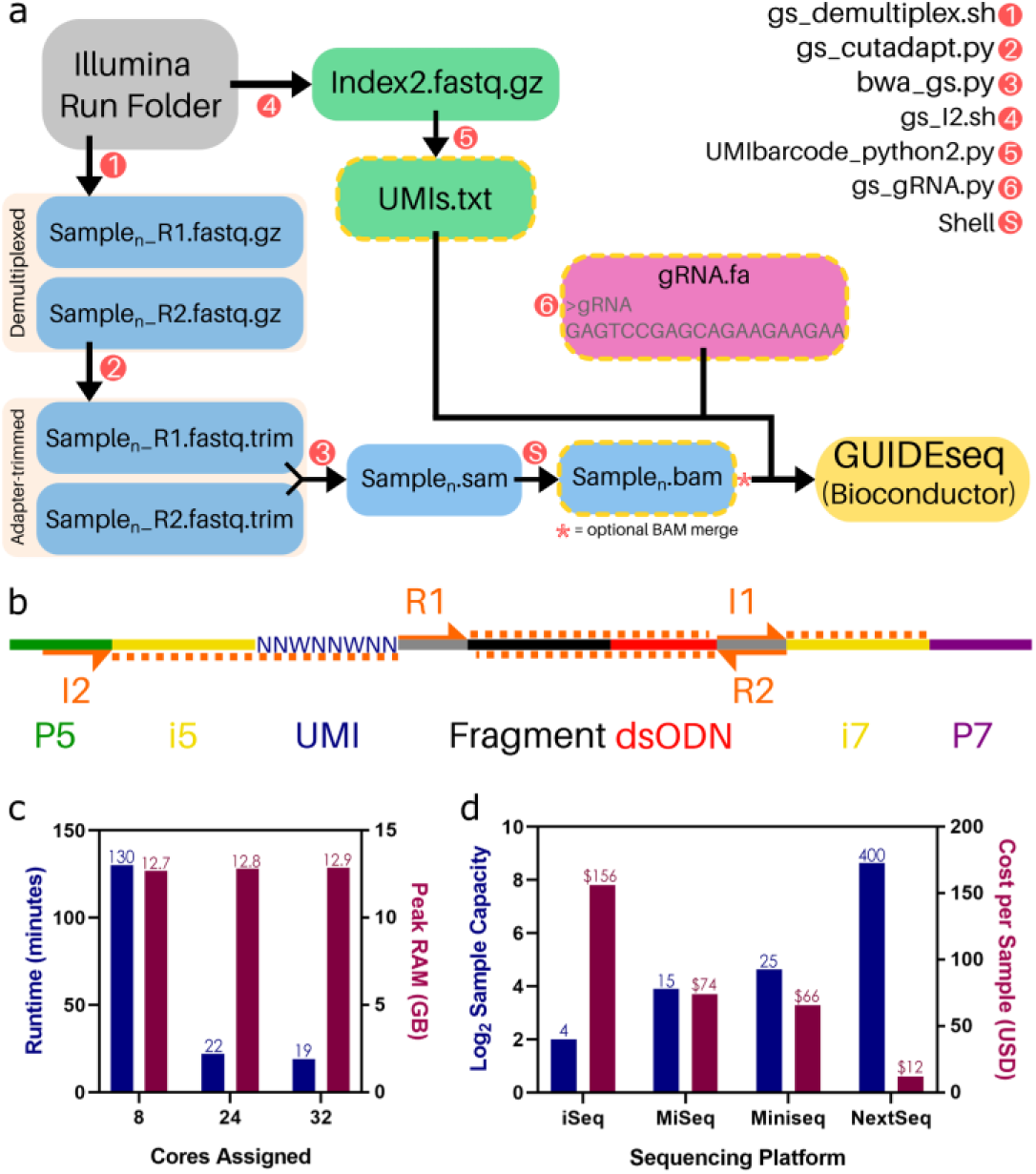
(a) GS-Preprocess workflow (b) Amplicon and Index read orientation shown on a standard Illumina-compatible GUIDE-seq library. Adapted from Tsai et al. 2014 (c) GS-Preprocess memory requirements and runtime for a 10^7^-read library pool across three degrees of multithreading. (d) Sequencing cost per GUIDE-seq library (10^6^ reads allocated to combined plus- and minus-strands) across common Illumina sequencers assuming highest-throughput, 150-cycle cartridge is used.

Similar to GUIDEseq BC, the Aryee Lab GUIDE-seq analysis pipeline is easy-to-use and well-explained. Users who wish to use GS-Preprocess output from a non-Miseq run for off-target profiling with https://github.com/aryeelab/guideseq#identify can do so by appending UMI references to BAMs.

### Benchmarking

To determine the resource allocation best fit for our software, we benchmarked GS-Preprocess runtime and memory consumption (RAM usage) across 8, 24, and 32 cores to process a 10^7^-read, 10-sample GUIDE-seq library pool (Fig. 1c). We found that the preprocessing pipeline can be completed on virtually any computing cluster or high-powered local machine in ∼20 min.

Given that our pipeline enables GUIDE-seq compatibility with sequencers that allow for increased multiplexing, we wanted to quantify its impact on method time and cost. We calculated the cost and maximum sample plexity for common Illumina sequencing platforms (Fig. 1d), assuming 10^6^ reads allocated to each sample (i.e., 500,000 plus- and minus-strand reads) run on a high-output, 300-cycle reagent cartridge purchased at list price (Illumina). Along with reductions in runtime and batch number, we observed a considerable decrease in cost per sample with increased multiplexing.

### Advantages over existing pipelines

GS-Preprocess integrates Illumina’s basecalling and demultiplexing software (bcl2fastq) and uses a standard CSV-format Illumina Sample Sheet to format GUIDE-seq libraries sequenced on *any* Illumina platform and *without* sequencer pre-configuration. By containerizing our software with docker, we have eliminated dependency installation steps. This may be particularly helpful for those unable to complete steps listed here: https://github.com/aryeelab/guideseq#dependencies), and assures long-term usability that is resistant to updates in Illumina sequencers or other software dependencies. This expands the usability of GUIDE-seq from labs with a configurable, in-house MiSeq to virtually any investigator with access to institutional or commercial NGS services. By allowing users to sequence on higher-throughput machines (e.g. NextSeq), GS-Preprocess increases sample plexity from dozens (with MiSeq) to several hundred, substantially reducing time and cost associated with large-scale off-target profiling.

## Discussion

GS-Preprocess is an-open source, dockerized package that generates all files necessary for off-target analysis of GUIDE-seq libraries in a single line of input code. This pipeline overcomes previous sequencer/multiplexing restrictions and eliminates the need for a YAML sample manifest. In doing so, GS-Preprocess makes GUIDE-seq a more accessible, user-friendly, and cost-effective method for the increasing number of research groups utilizing CRISPR-Cas technologies.

## End Matter

### Author Contributions and Notes

TCR, NA and PL designed read preprocessing steps. TCR, HEP and LJZ wrote software. TCR and HEP built docker containers. TCR and LJZ authored this manuscript.

The authors declare no conflict of interest.

This article contains supporting information at https://github.com/umasstr/gs-preprocess and https://hub.docker.com/r/umasstr/gsp

## Acknowledgments

The authors would like to acknowledge Emily Mohn for constructive edits on this manuscript. The authors thank the Aryee for its impeccably maintained Github repository, aryeelab/guideseq.

